# Grb2 Phosphorylation Antagonizes EGFR-driven Ras Activation by Interfering with Condensate Assembly

**DOI:** 10.1101/2024.09.05.611544

**Authors:** Henry T. Phan, Chun-Wei Lin, Brittany L. Stinger, Joseph B. DeGrandchamp, L.J. Nugent Lew, Serena J. Huang, Jay T. Groves

## Abstract

Upon ligand binding, the kinase domain of EGFR phosphorylates multiple tyrosine residues on the receptor cytoplasmic tail through a trans-autophosphorylation process. Phosphotyrosine sites on activated receptors recruit Grb2, which further recruits SOS to initiate downstream signaling by activating Ras. Multivalent binding between SOS and Grb2, as well as direct Grb2:Grb2 interactions, contribute to formation of a protein condensate of activated EGFR. The condensed state of EGFR facilitates autoinhibition release in SOS and exerts regulatory control over signal propagation from activated EGFR to Ras. While kinase activity of EGFR is an essential driver of this signaling process, phosphorylation at residue Y160 on Grb2 blocks Grb2:Grb2 binding and can interfere with EGFR condensation. Here, using a reconstituted system, we examine how titrating kinase activity in the EGFR system can both promote and inhibit signal output to Ras. The results reveal how effects of tyrosine kinase inhibition can, under some circumstances, promote Ras activation by inhibiting negative feedback through Grb2 phosphorylation and disruption of a Grb2 SH2/SH3 dimer interface.

**Statement of Significance:** Activated EGFR forms a biomolecular condensate, via linkage of multiple EGFR through Grb2 and SOS, and the condensation state of EGFR influences signal propagation to Ras. While tyrosine phosphorylation is a critical step in EGFR activation, phosphorylation of Grb2 can have an inhibitory effect on EGFR condensation and subsequent Ras activation. Under some conditions, kinase inhibition can promote signaling from EGFR to Ras.

## Introduction

Epidermal growth factor receptor (EGFR) is an important receptor tyrosine kinase, which is also the target of many therapeutic drugs. EGFR controls key cellular functions, including growth and proliferation, by signaling to Ras and the mitogen-activated protein kinase (MAPK) pathway. The basic molecular level features of the EGFR signaling mechanism are fairly well understood (1). In contrast, recent discovery of EGFR participation in a two-dimensional protein condensate on the membrane (2)—closely resembling the LAT protein condensate discovered in T cells (3–6)—is opening a wide range of new concepts in how EGFR signaling might be regulated.

EGFR consists of an extracellular ligand-binding module, a single transmembrane helix, a cytoplasmic tyrosine kinase domain, and an unstructured C-terminal tail (7). Under non-stimulating conditions, EGFR is autoinhibited and primarily monomeric. Upon addition of a ligand, EGFR dimerizes, and the kinase domain of one receptor phosphorylates tyrosine residues on the C-terminal tail of the other (1). In particular, phosphorylation of residues Y1068, Y1086, Y1148, and Y1173 recruit growth factor receptor-bound protein 2 (Grb2) through the Grb2 Src Homology 2 (SH2) domain (8). Grb2 then recruits the guanine nucleotide exchange factor Son of Sevenless (SOS) to the membrane (5, 9) through interaction between the SOS proline-rich domain and Grb2 Src Homology 3 (SH3) domains (10). SOS is recruited in an autoinhibited state and undergoes an autoinhibition release process at the membrane prior to initiating Ras activation. Multivalency in both EGFR:Grb2 binding (8) and in Grb2:SOS binding (10), as well as direct Grb2:Grb2 interactions (11–15) enable activated EGFR to form a protein condensate through a bond percolation network (2).

Although not as well studied as LAT condensates in T cells, the EGFR condensate shares many of the same molecules and physical properties. In particular, Grb2 and SOS are dominant players in the crosslinking interactions that enable condensation in both systems. Additionally, efficient release of autoinhibition in SOS, and subsequent Ras activation, require the condensed state in both systems (2, 4, 5). The Grb2:SOS:Grb2 linkage was the first studied in reconstitution, but it was later found that Grb2 alone is sufficient to drive condensation of both EGFR and LAT (2). Grb2-mediated condensation in reconstitution experiments relies on a Grb2:Grb2 interaction that occurs between Grb2 molecules bound to phosphorylated tyrosine residues. Various Grb2 dimeric interfaces have been observed in crystallographic studies (11–15), one of which contains an interaction interface between the C-SH3 domain of one Grb2 molecule and the SH2 domain of another Grb2 molecule (12). In this SH2/SH3 dimer, residue Y160 on one Grb2 molecule forms a hydrogen bond with residue E87 of the other protomer (13). Phosphorylation of Y160 breaks the Grb2:Grb2 interaction (16) and was also observed to inhibit EGFR condensation in reconstitution studies (2). While Grb2 may not be an ideal substrate for EGFR kinase (9, 17, 18), the close proximity of active kinase with Grb2 in condensates could strongly promote promiscuous phosphorylation. We hypothesize that Grb2 phosphorylation could introduce a negative regulatory coupling on the activation of Ras by EGFR through the dependency of SOS-Ras activity on condensation state.

Here, we describe experimental reconstitution of the C-terminal tail of EGFR (EGFR^TAIL^) on supported lipid bilayers (SLBs) along with detailed microscopic imaging studies of EGFR-driven activation of Ras. Kinase activity levels were titrated, either by varying kinase density on the membrane or using a tyrosine kinase inhibitor, while both EGFR condensation state and the rate of Ras activation (by Grb2-recruited full-length SOS) were monitored. Titrating up kinase activity initially leads to increased EGFR condensation and Ras activation. However, at higher kinase activity levels, the EGFR condensation state appears more dispersed, and Ras activation rates are correspondingly lower. Through a series of studies with Grb2 mutations, including a Y160E mutation that blocks the SH2/SH3 dimer, we confirm that Grb2 Y160 phosphorylation is the driver of reduced Ras activation at high kinase levels through its interference with EGFR condensation.

EGFR overactivity contributes to many cancers and tyrosine kinase inhibitors (TKIs) are regularly used as cancer therapeutics intended to block EGFR-driven Ras/MAPK signaling (19, 20). We observe that adding a kinase inhibitor to an EGFR system exhibiting kinase overactivity can drive increased Ras activation. The results described here illustrate how effects of Grb2 phosphorylation enable kinase activity to drive both positive and negative aspects of the EGFR-Ras/MAPK signaling.

## Materials and Methods

### Chemicals

1,2-dioleoyl-sn-glycero-3-phosphocholine (DOPC), 1,2-dioleoyl-sn-glycero-3-[(N-(5-amino-1-carboxypentyl)iminodiacetic acid)succinyl] (nickel salt) (Ni^2+^-NTA-DOGS), 1,2-dioleoyl-sn-glycero-3-phosphoethanolamine-N-[4-(p-maleimidomethyl)cyclohexane-carboxamide] (sodium salt) (PE MCC), and L-α-phosphatidylinositol-4,5-bisphosphate (Brain, Porcine) (ammonium salt) (Brain PIP_2_) were purchased from Avanti Polar Lipids. Alexa Fluor 488 and Alexa Fluor 555 dyes were purchased from ThermoFisher. SNAP-Surface Alexa Fluor 647 was purchased from New England Biolabs. Bovine Serum Albumin (BSA), 2-Mercaptoethanol (BME), NiCl_2_, H_2_SO_4_, ATP, GDP, GTP, phosphatase inhibitor cocktail 2 and 3, and Dasatinib were purchased from Sigma-Aldrich. H_2_O_2_ and Tris buffer saline (TBS) were purchased from Fisher Scientific. MgCl_2_ was from EMD Chemicals.

### Protein purification

**EGFR^TAIL^** EGFR C-terminal tail (residues 991 to 1186, not including the 24-residue signaling peptide) was purified as described previously (2) using an N-terminal His6 tag and SUMO tag with a Tobacco Etch Virus (TEV) cleaving site. The His6 tag was left intact for membrane tethering.

**LAT** Cytoplasmic LAT (residues 27-233, human) with mutation C117F was purified as described previously (21) using an N-terminal His6 tag with a TEV cleaving site. The His6 tag was left intact for membrane tethering.

**Grb2** Full length and mutant Grb2 were purified as described previously (2) using an N-terminal His6 tag. The N-terminal His6 tag was removed with TEV protease. Grb2 was further equilibrated at 37 °C for 10 minutes to reestablish equilibrium between monomer and dimer.

**Hck** Hck kinase domain (residues 83-526, human) was purified as described previously for Src-family kinases (22) using an N-terminal His6 tag and TEV cleaving site. The His6 tag was left intact for membrane tethering.

**SOS^PR^** The proline rich domain of SOS (residues 1051-1333, human) was expressed with an N-terminal His6-MBP-fusion tag with a TEV cleaving site. The protein was expressed and purified as described previously (4).

**SOS^FL^** Full-length SOS protein was prepared via a split intein approach. The N- and the C-terminal fragments of human SOS1 were expressed in BL21 (DE3) bacteria and purified separately. The two fragments with high purity were ligated by the intein reaction to generate SOS^FL^. The protein was purified as described previously (5).

**RBD-K65E** pETM11 vector containing the ORF, His6-GST-PreScission-SNAPtag-Raf1 RBD (residue 56-131, K65E) derived from the Raf-1 human gene was transformed into BL21 (DE3) bacteria. The N-terminal His6-GST tag was removed with PreScission before use. The protein was purified as described previously (5).

### Protein labeling

EGFR^TAIL^, LAT, and SOS^FL^ were diluted to 100 μM in 5 mM TCEP. Alexa Fluor 488 and 555 maleimide dyes (ThermoFisher) were dissolved to make 10-20 mM stock. The dyes were added to protein solutions and allowed to react for 1 hr at room temperature. The reaction was quenched with 10 mM dithiothreitol (DTT) for 10 min. Amicon Ultra centrifugal filters were used to wash away free dye and to concentrate protein. Protein was loaded into an analytical grade size exclusion column (SE, GE Healthcare) for further purification. Labeled fraction was determined by comparison of absorption of the dyes λ_max_ to the protein’s UV absorption at 280 nm corrected for the dye’s absorption at 280 nm.

RBD-K65E was labelled with SNAP-Surface Alexa Fluor 647 (S9136S, New England Biolabs) according to the manufacturer protocol. Excess dye was removed with gravity desalting columns (PD10, Cytica) and by SEC via FPLC (Superdex 75 increase, 24 mL bed volume, Cytiva).

### SUV and supported lipid bilayers

Small unilamellar vesicles (SUVs) were prepared using 1,2-dioleoyl-sn-glycero-3-phosphocholine (DOPC), 1,2-dioleoyl-sn-glycero-3-[(N-(5-amino-1-carboxypentyl)iminodiacetic acid)succinyl] (nickel salt) (Ni^2+^-NTA-DOGS), 1,2-dioleoyl-sn-glycero-3-phosphoethanolamine-N-[4-(p-maleimidomethyl)cyclohexane-carboxamide] (sodium salt) (PE MCC), and L-α-phosphatidylinositol-4,5-bisphosphate (Brain, Porcine) (ammonium salt) (Brain PIP_2_) (Avanti Polar Lipids). Purchased lipids were mixed into a piranha etched round bottom flask to 2 mg of total lipid in a ratio of 92:2:2:4 (by molar percent) of DOPC:PIP_2_:PE MCC:DGS-NTA(Ni). After mixing, lipids were dried using a rotary evaporator for 10 min at room temperature and an additional 3 min at 42 °C. Lipid films were further dried under continuous N_2_ flow for ∼15 min. Lipid films were rehydrated with 2 mL of milli-Q water, resulting in a lipid concentration of 1 mg/mL and vigorously vortexed to ensure lipid film liftoff. SUVs were made by sonicating the lipid mixture with a 3 mm stepped tip (Sonics VCX750 – 33% amplitude, 20 s sonication, 50 s rest, and 5 complete cycles). SUVs were stored at 4 °C and used within 1 week of preparation.

Precision Schott D 263 M glass coverslips (#1.5H, 75 x 25 mm, custom ordered from ThorLabs) were cleaned in 50% IPA:milli-Q water with bath sonication for 30 min. Glass substrates were then piranha etched with a 1:3 ratio of 30% H_2_O_2_ and concentrated H_2_SO_4_ for 7 min. Coverslips were washed excessively with milli-Q water after etching, stored in milli-Q water at room temperature, and used within 1 week. For use, etched coverslips were blow-dried under N_2_ and attached to a 6-channel slide (sticky-Slide VI 0.4, Ibidi). Chambers were cured at 37 °C for 30 min under heavy weight.

SLBs were formed by rupture of SUVs. 1 mg/mL SUV solution was mixed with 10 mM TBS (1:1 ratio by volume), and 200 μL of the diluted solution was injected into the chamber with a 30 min incubation time. Channels were washed with 10 mM TBS, and 100 mM NiCl_2_ was added to each channel to reload Ni^2+^-NTA-DOGS for 5 min. Channels were washed and then blocked with 1 mg/mL BSA in TBS buffer for 10 min. After washing, H-Ras was anchored on PE MCC through maleimide chemistry by incubating 0.5 mg/mL H-Ras in the chambers for 2.5 hr. Free PE MCC was quenched with 5 mM BME for 20 min. EGFR^TAIL^ and Hck were anchored on Ni^2+^-NTA-DOGS through His-tag and Ni^2+^ NTA chemistry and through incubation at 50 nM EGFR^TAIL^ for condensation experiments (Figure 1, Figure 2, Figure 3, Figure 5, Figure 5, Figure S1) or 20 nM EGFR^TAIL^ for Ras activation experiments (Figure 6, Figure S4, Figure S5, Figure S6) for 45 min. LAT experiments were performed at 150 nM LAT (Figure S1). During this step, 100 μM GDP was included to ensure Ras was loaded with GDP. After washing, EGFR^TAIL^ was phosphorylated by incubating 1 mM ATP and 5 mM MgCl_2_ in TBS for 20 min. The incubation period also allows for weakly chelated proteins to dissociate. The mobility of the proteins was examined using fluorescence recovery after photobleaching (FRAP).

**Figure 1:**
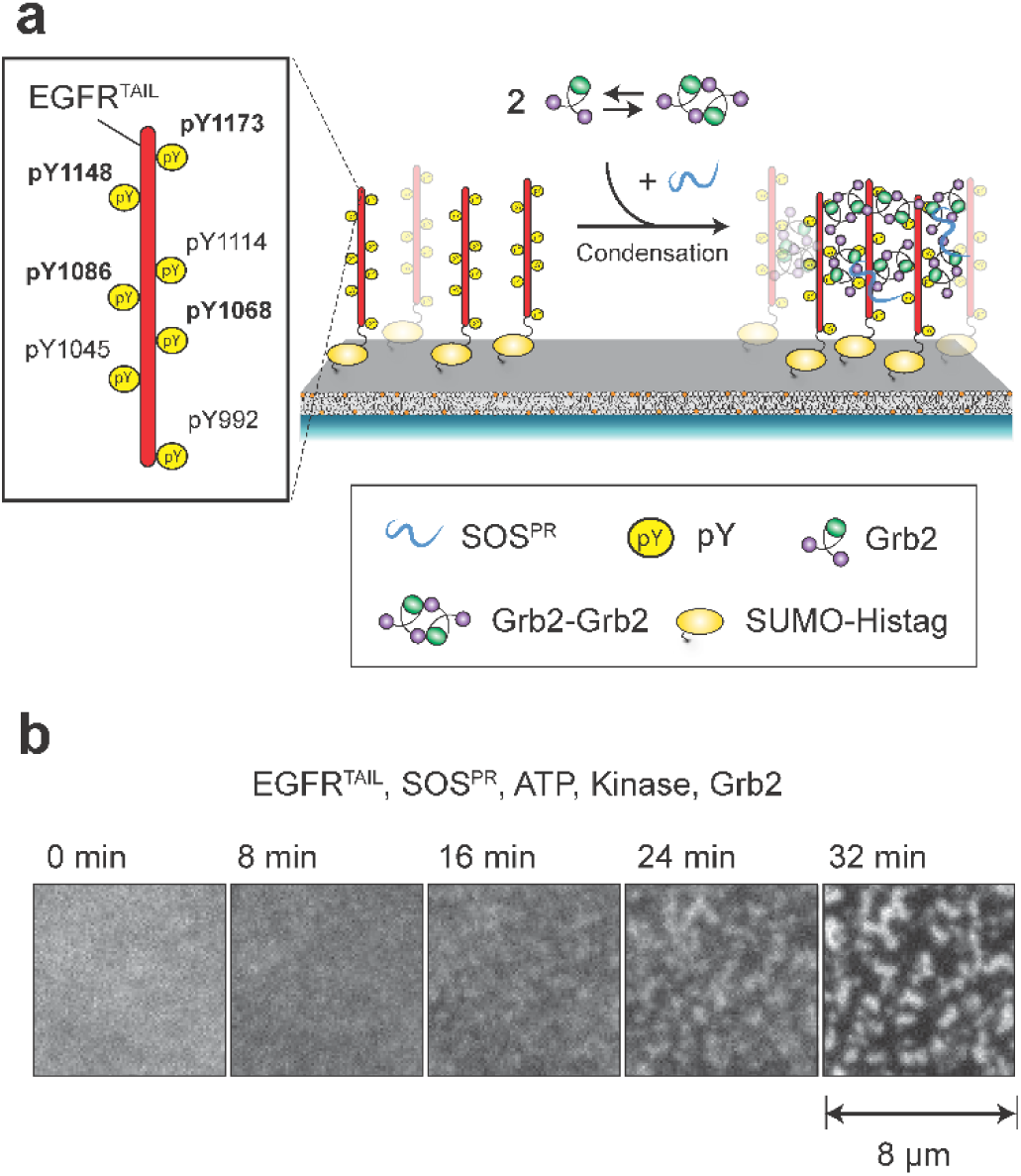
EGFR^TAIL^ reconstituted on supported lipid bilayers (SLBs) undergoes protein condensation upon addition of Grb2 and SOS^PR^. (A) Schematic of the reconstitution experiment. EGFR^TAIL^ can be phosphorylated on several tyrosine sites, of which Y1068, Y1086, Y1148, and Y1173 are implicated in Grb2 binding. Grb2 dimerizes through a Grb2-Grb2 interface and additionally interacts with the PR domain of SOS (SOS^PR^). Grb2 and SOS^PR^ are injected into the chamber and are recruited to EGFR^TAIL^. The crosslinking between EGFR^TAIL^:Grb2:SOS^PR^ drives the assembly of a protein condensate. EGFR^TAIL^ is initially in a dispersed phase (Left), but the addition of Grb2 and SOS^PR^ drives EGFR^TAIL^ into a condensed phase (Right). (B) TIRF images of EGFR^TAIL^-AF488 on SLBs. After the addition of Grb2 and SOS^PR^, EGFR^TAIL^ condenses over the course of 32 min.

**Figure 2:**
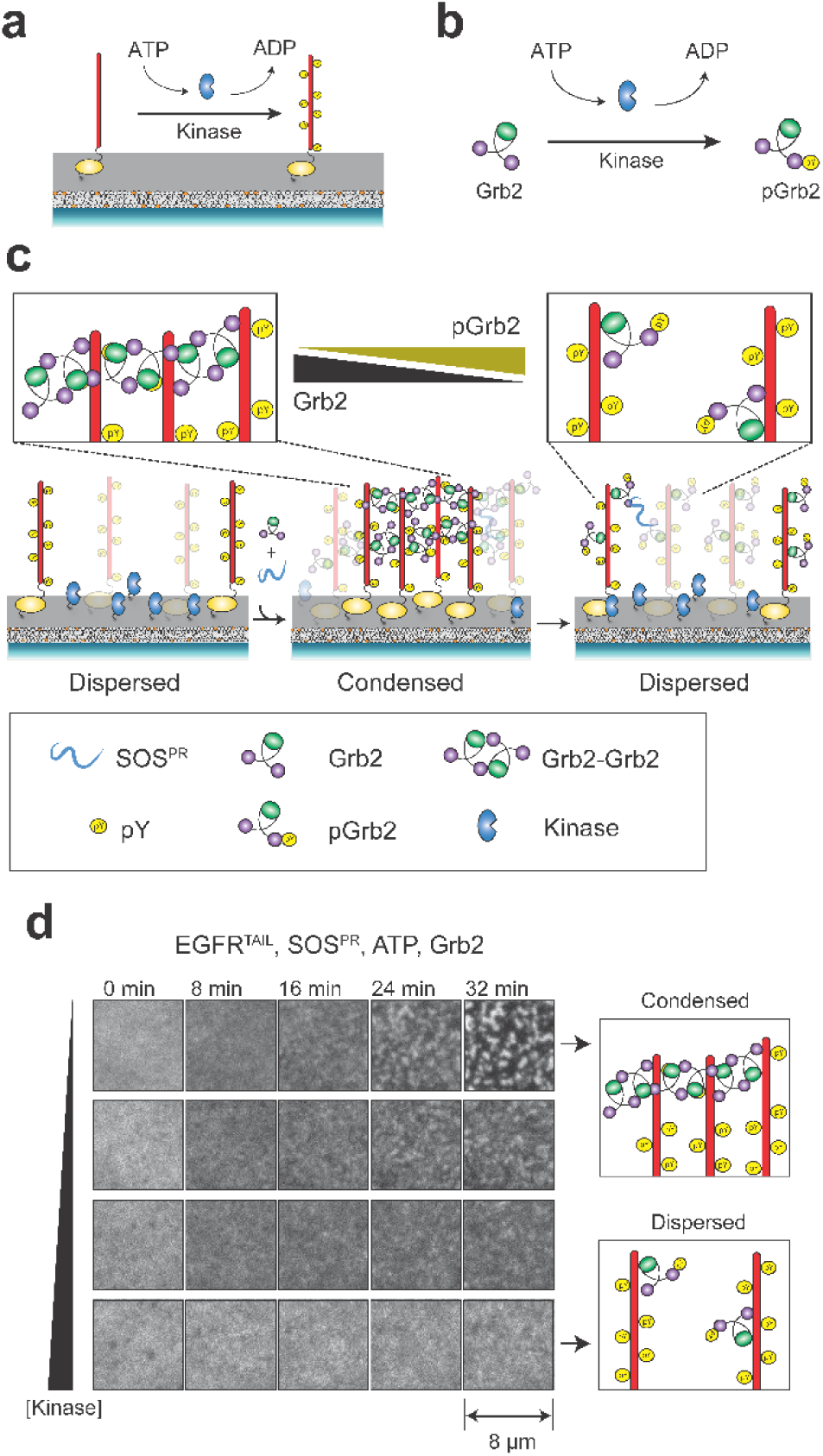
Phosphorylation of Grb2 inhibits EGFR condensation. (A) Schematic of the role of kinases in EGFR^TAIL^ phosphorylation. Kinases phosphorylate EGFR^TAIL^ on several tyrosine residues which recruit Grb2. (B) Schematic of the role of kinases in Grb2 phosphorylation. Kinases phosphorylate Grb2 to drive monomer formation. (C) Schematic of the role of kinases in EGFR^TAIL^ condensation. Before the addition of Grb2 and SOS^PR^, EGFR^TAIL^ is in a dispersed state (Left). After the addition of Grb2 and SOS^PR^, EGFR^TAIL^ condenses with Grb2 and SOS^PR^ acting as crosslinkers (Middle). At later time points, kinases phosphorylate Grb2, breaking dimer formation. EGFR^TAIL^ condensation is dissolved by the lack of Grb2 dimers because crosslinking through Grb2:SOS^PR^ is insufficient (Right). (D) TIRF images of EGFR^TAIL^-AF488 on SLBs functionalized with increasing incubation kinase concentrations of 10 nM, 50 nM, 100 nM, and 150 nM. After the addition of Grb2 and SOS^PR^, EGFR^TAIL^ condenses at low kinase conditions (Top) but remains homogenous at high kinase conditions (Bottom). Hck was utilized to phosphorylate both EGFR^TAIL^ and Grb2.

**Figure 3:**
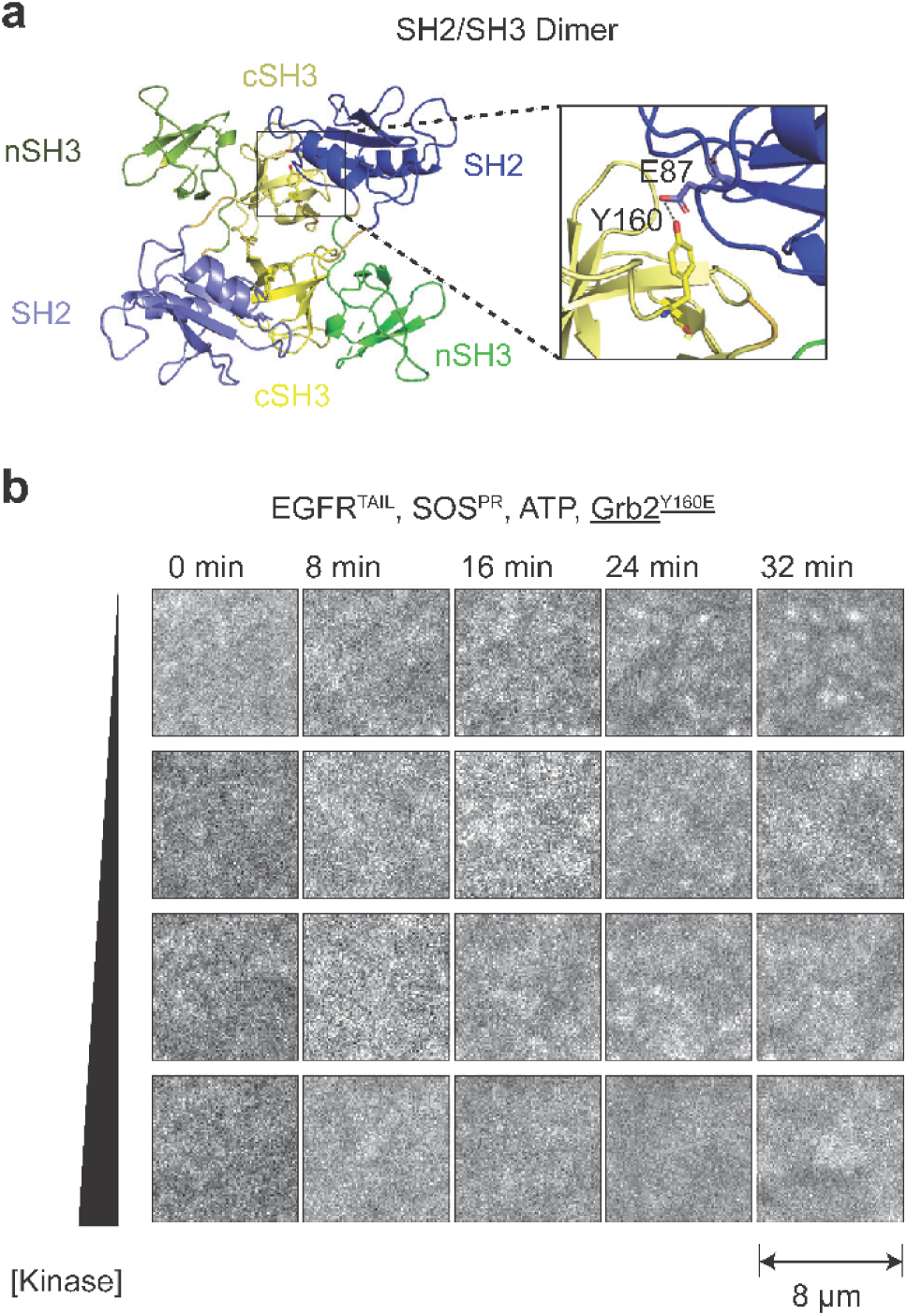
Breakage of the Grb2 SH2/SH3 dimer inhibits EGFR condensation (A) Crystal structure of the Grb2 SH2/SH3 dimer (PDB:1GRI) (13). The interaction between Y160 and E87 is magnified. (B) TIRF images of EGFR^TAIL^-AF488 on SLBs functionalized with increasing incubation kinase concentrations of 10 nM, 50 nM, 100 nM, and 150 nM. After the addition of Grb2^Y160E^ and SOS^PR^, EGFR^TAIL^ remains dispersed at all kinase concentrations.

**Figure 4:**
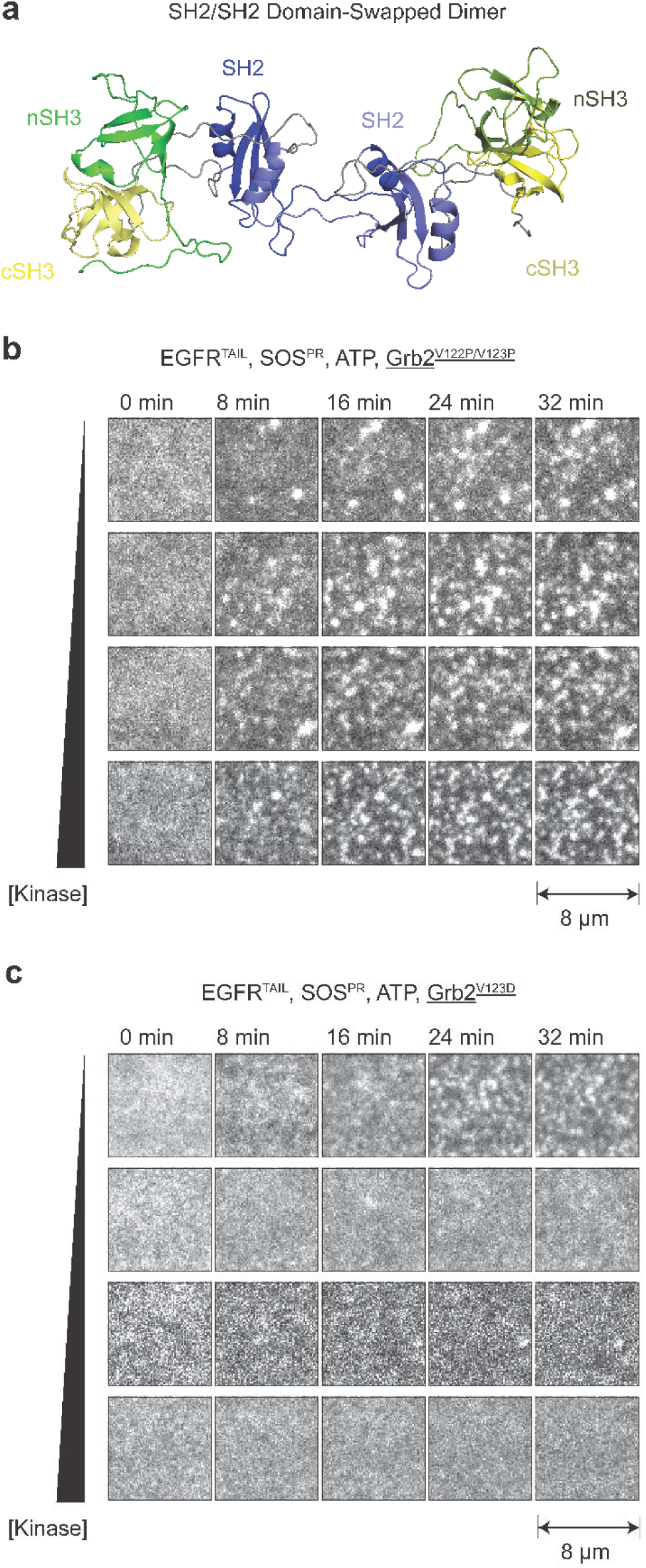
Stabilization of a proposed Grb2 SH2/SH2 domain-swapped dimer can promote EGFR condensation (A) Proposed model of the Grb2 SH2/SH2 domain-swapped dimer determined from SEC-MALS-SAXS (SASPD78) (11). (B) TIRF images of EGFR^TAIL^-AF488 on SLBs functionalized with increasing incubation kinase concentrations of 10 nM, 50 nM, 100 nM, and 150 nM. After the addition of Grb2^V122P/V123P^ (mutant suggested to stabilize a Grb2 domain-swapped dimer) and SOS^PR^, EGFR^TAIL^ condenses at all kinase concentrations. Hck was utilized to phosphorylate both EGFR^TAIL^ and Grb2. (C). TIRF images of EGFR^TAIL^-AF488 on SLBs functionalized with increasing incubation kinase concentrations of 10 nM, 50 nM, 100 nM, and 150 nM. After the addition of Grb2^V123D^ and SOS^PR^, EGFR^TAIL^ condenses at low kinase conditions (Top) but remains homogenous at high kinase conditions (Bottom). Hck was utilized to phosphorylate both EGFR^TAIL^ and Grb2.

**Figure 5:**
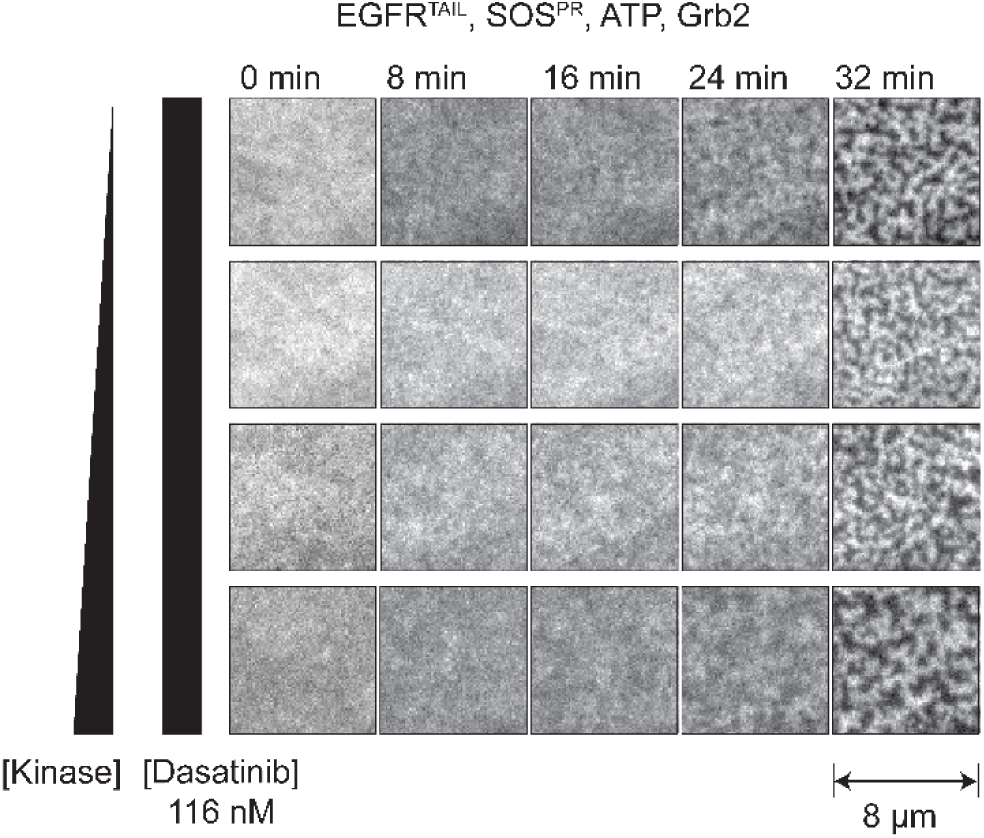
Tyrosine kinase inhibition promotes EGFR condensation. TIRF images of EGFR^TAIL^-AF488 reconstituted on SLBs with increasing incubation kinase concentrations of 10 nM, 50 nM, 100 nM, and 150 nM. Tyrosine kinase inhibitor is added along with Grb2 and SOS^PR^. EGFR^TAIL^ with kinase inhibitor condenses at all kinase concentrations.

**Figure 6:**
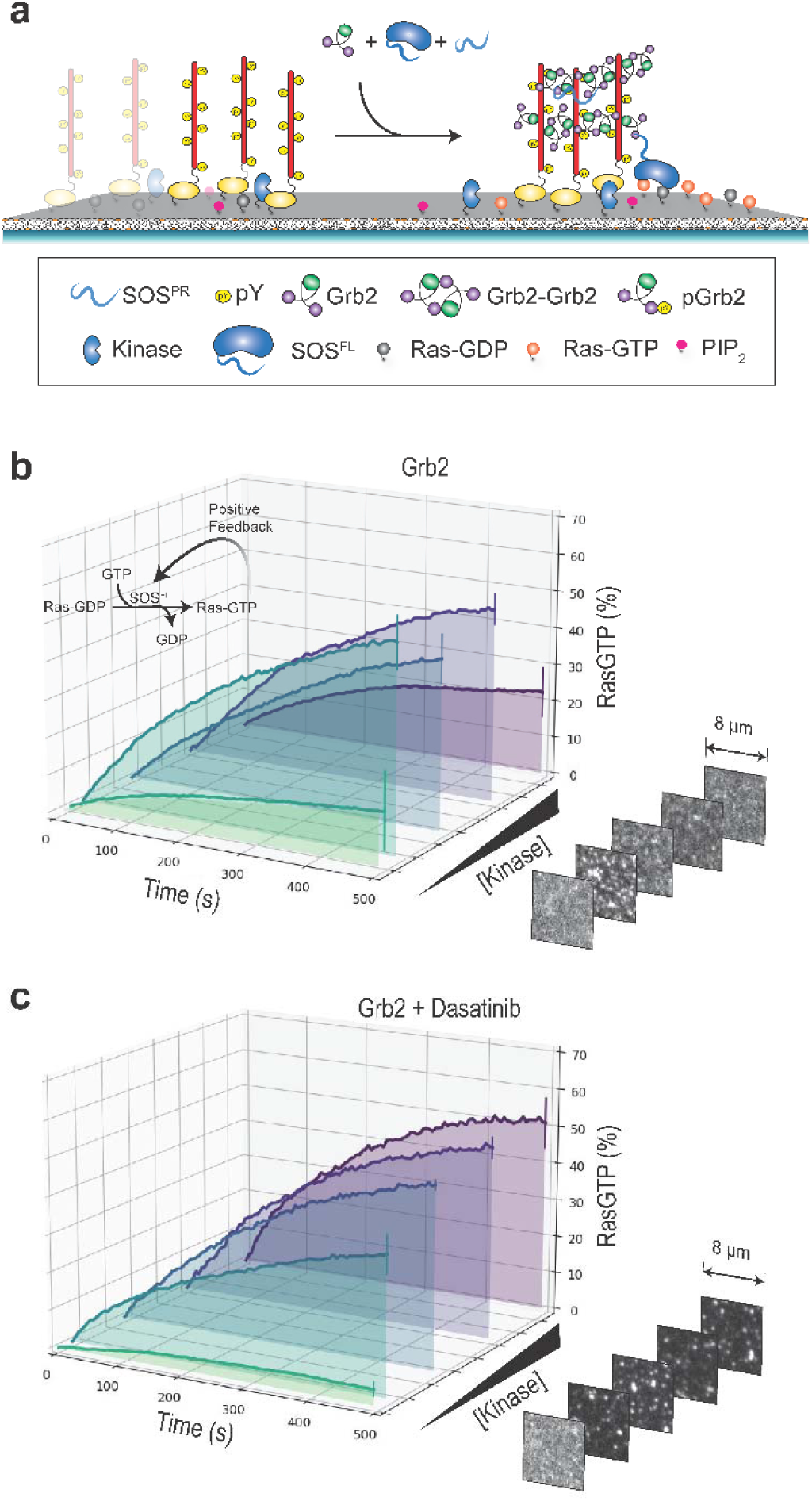
Tyrosine kinase inhibition can promote Ras activation through a double negative effect. (A) Schematic of Ras activation by EGFR^TAIL^, Grb2, SOS^PR^, and SOS^FL^ reconstituted on SLBs. Grb2, SOS^PR^, SOS^FL^-AF555, RBD-AF647, and GTP (1 mM) were injected onto SLBs functionalized with EGFR^TAIL^-AF488, Hck, and Ras-GDP. Grb2 recruits SOS^FL^ to the bilayer which activates Ras by converting Ras-GDP to Ras-GTP. Activated Ras is measured by the fluorescence intensity of RBD-AF647, which specifically binds to Ras-GTP. (B) Mean kinetic traces of ≥ 2 replicates of Ras activation with increasing incubation kinase concentrations of 0 nM, 10 nM, 50 nM, 100 nM, and 150 nM. Error bars represent the standard error of the mean (SEM) of ≥ 2 replicates. Displayed are representative TIRF images of EGFR^TAIL^-AF488 on Ras-functionalized bilayers after the addition of Grb2, SOS^PR^, and SOS^FL^-AF555. High kinase concentrations inhibit Ras activation and correspond to homogenous distributions of EGFR^TAIL^. (C) Mean kinetic traces of ≥ 2 replicates of Ras activation with increasing incubation kinase concentrations of 0 nM, 10 nM, 50 nM, 100 nM, and 150 nM and with the addition of kinase inhibitor. Error bars represent the SEM of ≥ 2 replicates. Displayed are representative TIRF images of EGFR^TAIL^ on Ras-functionalized bilayers after the addition of Grb2, SOS^PR^, SOS^FL^-AF555, and kinase inhibitor. Kinase inhibition rescues Ras activation at high kinase concentrations and correspond to a condensed state of EGFR^TAIL^.

### Total internal reflection fluorescence (TIRF) microscopy

Imaging experiments utilizing TIRF were performed on an inverted Nikon Eclipse Ti microscope with an 100x Nikon oil-immersion TIRF objective (1.49 NA). A 1.5x lens tube (Nikon) was used for a final magnification of 150x. A motor-controlled stage (MS-2000, ASI) was used for positioning, and images were acquired using an iXon Ultra 897 EMCCD camera (Andor Technology). The laser unit has diode lasers of 488 nm, 561 nm, and 637 nm (OBIS laser diode, Coherent) controlled with a custom laser driver (Solamere Technology). The laser was collimated with the objective at the same approximate z-focus between experiments. A single quad-band dichroic cube was used for all acquisitions (ZT405/488/561/640rpcv, Chroma) and additionally filtered with an ET600/50M filter (Semrock). The microscopy and accessory hardware were controlled with Micro-Manager version 4.0 (23). All experiments were conducted at 21-23 °C.

### Condensation Imaging

After EGFR^TAIL^ phosphorylation, 1.5 μM Grb2, 188 nM SOS^PR^, 0.1 mg/mL BSA in TBS, and 1 mM ATP were added into the chamber as a 100 μL injection. 116 nM Dasatinib (Sigma-Aldrich) was added in the kinase inhibitor studies (Figure 5). Images were acquired with a 20 ms exposure time and 250 EM gain. EGFR^TAIL^ was excited with a 488 nm diode laser (OBIS laser diode, Coherent) with 1 mW (∼0.25 mW at objective) of power. Images were collected at 1 min intervals with acquisition durations as long as 1 hr. Image analysis and contrast adjustments were made in Fiji (24).

### Ras Activation Imaging

Ras activation studies were performed similarly as described previously (2). Briefly, to initiate downstream signaling of Ras, 1.5 μM Grb2, 186 nM SOS^PR^, 2 nM SOS^FL^-AF555, 50 nM RBD-AF647, 0.1 mg/mL BSA in TBS, 1 mM GTP, and 1 mM ATP were added into the chamber as a 100 μL injection. Concentrations between 1.16 nM and 232 μM of Dasatinib (Sigma-Aldrich) (Figure 6, Figure S4) were added in the kinase inhibitor studies. SOS^FL^ is recruited to the membrane by Grb2 and begins to activate. Once activated, SOS^FL^ will processively convert Ras-GDP to Ras-GTP. RBD-647 binds specifically to Ras-GTP (5), and activated Ras was measured by the fluorescence intensity of RBD-647 on TIRF images. At the end of the Ras trace, 30 nM SOS^CAT^, which is the catalytic domain of SOS^FL^, was used to fully activate Ras in the SLB. After 20 min, the average RBD intensity was used as a normalization factor for comparison of different chambers with various Ras densities due to random variations from different experiment days. The concentrations of EGFR, Grb2, and SOS^FL^ were determined to make the time scale of Ras activation about 10 min. Images were acquired with a 100 ms, 100 ms, and 20 ms exposure time and 100 EM gain, 500 EM gain, and 500 EM gain for 488 nm, 561 nm, and 637 nm channels respectively. 488 nm, 561 nm, and 637 nm lasers were at 1 mW, 2 mW, and 3 mW power respectively. Images were collected at 5 sec intervals for 25 min. Quantitative image analysis was done in Python.

## Results

### Reconstitution of the EGFR condensate on supported bilayers

The EGFR condensate was reconstituted on SLBs from purified proteins following established protocols (2). Briefly, the C-terminal tail of EGFR (residues 991 to 1186, not including the 24-residue signaling peptide) (EGFR^TAIL^) was expressed with a His6 tag and a small ubiquitin-related modifier (SUMO) fusion tag at the N-terminus of the construct. This EGFR^TAIL^ construct was labeled with Alexa Fluor 488 (66% efficiency) and anchored onto SLBs through Ni^2+^-histidine chelation with 1,2-dioleoyl-sn-glycero-3-[(N-(5-amino-1-carboxypentyl)iminodiacetic acid)succinyl] (nickel salt) (Ni^2+^-NTA-DOGS) lipids. SLBs are prepared with 4% Ni^2+^-NTA-DOGS by mole percent with 1,2-dioleoyl-sn-glycero-3-phosphocholine (DOPC) as the main lipid constituent. The SUMO tag acts both as a solubility tag for increased purification yield and as a spacer that mimics the area of the EGFR kinase domain. To phosphorylate EGFR^TAIL^, Src-family kinase Hck was attached to SLBs through Ni^2+^-histidine chelation. Since EGFR is not efficiently phosphorylated by Src-family kinases (25), a prephosphorylation strategy was employed in which EGFR^TAIL^, Hck, and adenosine triphosphate (ATP) are incubated together before the addition of any adaptor proteins. Addition of Grb2 can induce condensation of the phosphorylated EGFR^TAIL^. However, condensation can be achieved at lower concentrations when small amounts of the SOS proline-rich domain (SOS^PR^) are included as well (Figure 1A). Total internal reflection fluorescence (TIRF) images confirm that before the addition of Grb2 and SOS^PR^, EGFR^TAIL^ is homogeneously distributed on the SLB. Upon injection of adaptor proteins, EGFR^TAIL^ undergoes a protein-condensation phase transition and segregates into highly dense (bright) and disperse (dark) regions of EGFR^TAIL^ (Figure 1B).

### Kinase-driven phosphorylation of Grb2 antagonizes EGFR condensation

EGFR^TAIL^ phosphorylation and condensation is dependent on kinase activity. However, crosslinking interactions between Grb2 molecules via the SH2/SH3 Grb2 dimer interaction, which contribute to EGFR^TAIL^ condensation, can be broken by kinase activity. This Grb2 phosphorylation effect adds an antagonistic regulatory coupling that complicates how the system responds to kinase pressure. Figure 2A-C illustrates schematically how increasing kinase activity can initially drive condensation while leading to dispersed states at the highest kinase activity levels. We examine this effect by experimentally titrating up kinase activity by incubating the system with higher concentrations of Hck. At low levels of kinase activity, EGFR^TAIL^ condenses (Figure 2D, Top) after the addition of Grb2 and SOS^PR^. However, at high levels of kinase activity, EGFR^TAIL^ remains macroscopically homogenous (Figure 2D, Bottom). Experiments with LAT demonstrate the same kinase-driven negative regulatory mechanism (Figure S1). We confirm Grb2 phosphorylation through mass spectrometry (Table S1). We also confirm that Grb2 phosphorylation does not interfere with SOS recruitment (Figure S2).

### EGFR condensation is controlled by Grb2 SH2/SH3 dimer formation

Two distinct Grb2 dimer interfaces have been identified from crystallographic studies. The SH2/SH3 dimer interface involves a critical hydrogen bond between residue Y160 on one Grb2 molecule with residue E87 of the other protomer (13). We have confirmed that phosphorylation of Grb2 at Y160 disrupts EGFR^TAIL^ condensation, suggesting this interface is responsible for Grb2 crosslinking in these experiments. However, another dimeric interface dubbed the SH2/SH2 domain-swapped dimer has been identified, in which the α-helix of one Grb2 SH2 domain swaps with the α-helix of the SH2 domain of the other Grb2 molecule (11, 14, 15).

To further investigate potential roles of these different Grb2 dimer interfaces in EGFR condensation, we generated Grb2 mutants that stabilize or disrupt different Grb2 dimer conformations. Grb2 mutation Y160E (Grb2^Y160E^) lies within the SH2/SH3 dimer interface (Figure 3A) and prevents dimer formation (16). We confirm experimentally that when Grb2^Y160E^ and SOS^PR^ are added, EGFR^TAIL^ remains in a dispersed state regardless of kinase activity level (Figure 3B). In the Grb2 domain-swapped dimer structure, residues V122 and V123 lie on the hinge region (15) (Figure S3), and Grb2 mutations V122P and V123P (Grb2^V122P/V123P^) promote formation of the domain-swapped dimer (11) (Figure 4A). In experiments in which Grb2^V122P/V123P^ and SOS^PR^ were added to EGFR^TAIL^, we observed the system to condense at all kinase concentrations (Figure 4B). This confirms that Grb2^V122P/V123P^ promotes EGFR^TAIL^ condensation regardless of Grb2 phosphorylation state.

Lastly, we examined the Grb2 mutation V123D (Grb2^V123D^), which disrupts domain-swapped dimer formation (11) but may still form the SH2/SH3 dimer. In experiments performed with Grb2^V123D^ and SOS^PR^, EGFR^TAIL^ behaved similarly to corresponding experiments with wild-type Grb2. For both Grb2^V123D^ and wild type Grb2, EGFR^TAIL^ condensed at low levels of kinase activity (Figure 4C, Top) but remains dispersed at high levels of kinase activity (Figure 4C, Bottom). This data, taken together with the Grb2^Y160E^ data, suggests that Grb2 primarily forms the SH2/SH3 dimer in reconstitution and that the SH2/SH2 domain-swapped dimer is not a major contributing factor in these experiments.

### Tyrosine kinase inhibition can promote EGFR condensation

Tyrosine kinase inhibitors, which are used to reduce EGFR driven signaling activity, may also inhibit the negative regulatory effects of Grb2 phosphorylation on EGFR condensation. To examine this possibility, we studied effects of the Src-family specific kinase inhibitor Dasatinib (26, 27) on EGFR condensation under varying degrees of kinase pressure. In these experiments, Dasatinib is injected after the phosphorylation of EGFR^TAIL^, thus it will not interfere with Grb2 recruitment but will block Grb2 subsequent phosphorylation. In the presence of Dasatinib (116 nM), we observe EGFR^TAIL^ to form protein condensates over the entire range of kinase activity levels (Figure 5). This behavior differs substantially from inhibitor-free experiments, where we observed dispersed EGFR^TAIL^ distributions due to Grb2 phosphorylation at all but the lowest kinase levels (see Figure 2D). Similarly, Dasatinib promotes LAT condensation even under high concentrations of kinase (Figure S1). Although higher kinase activity levels negatively regulate EGFR^TAIL^ condensation, kinase inhibitors can negatively regulate this inhibitory mechanism.

### EGFR-mediated Ras activation exhibits nonmonotonic dependence on kinase activity

We next quantify how these effects on EGFR condensation state translate into rates of Ras activation by SOS. For these studies, we reconstitute the EGFR:Grb2:SOS:Ras signaling pathway as depicted schematically in Figure 6A. EGFR^TAIL^ along with the Src kinase, Hck, are anchored onto SLBs containing 2 mole % PIP_2_, which facilitates SOS^FL^ autoinhibition release and Ras activation. RasGDP is attached to the SLBs through cysteine-maleimide chemistry with 1,2-dioleoyl-sn-glycero-3-phosphoethanolamine-N-[4-(p-maleimidomethyl)cyclohexane-carboxamide] (sodium salt) (PE MCC) lipids (5, 28, 29). The reaction is initiated by adding GTP 1 mM, Grb2 1.5 μM, SOS^PR^ 186 nM, and fluorescently labeled SOS^FL^ 2 nM to the solution phase above the SLB. Grb2 binds to the phosphorylated tyrosine residues of EGFR^TAIL^ and subsequently recruits SOS^PR^ and SOS^FL^. Grb2:SOS:Grb2 as well as Grb2:Grb2 interactions drive crosslinking and condensation of EGFR^TAIL^ on the membrane surface. SOS^FL^ is initially recruited in an autoinhibited state and undergoes an autoinhibition release process on the membrane prior to beginning to catalyze nucleotide exchange in Ras. Ras activation from its GDP- to its GTP-bound state is monitored in real time using a modified Ras binding domain (RBD) derived from Raf (5) and fluorescently labeled with Alexa Fluor 647. This modified RBD selectively binds RasGTP with fast binding kinetics and provides an accurate readout on Ras activity levels (30). We then utilize TIRF imaging of RBD and EGFR^TAIL^ at the membrane to simultaneously visualize EGFR^TAIL^ condensation state and measure the rate of Ras activation.

Kinetic traces of Ras activation under a range kinase activity levels, with and without kinase inhibition, are plotted in Figure 6B and 6C. Without kinase inhibition, Ras activity levels exhibit a nonmonotonic dependence on kinase activity, with notably reduced rates of Ras activation at the highest kinase levels (Figure 6B). In contrast, in the presence of kinase inhibition by Dasatinib, progressively higher rates of Ras activation were observed with increasing kinase at all levels (Figure 6C). The corresponding state of EGFR^TAIL^ condensation under each condition is shown in the image stacks along the right side of Figure 6B and 6C. The rate of Ras activation shows a strong correlation with the condensation state, including with the Grb2 mutants studied above (Figure S5, Figure S6). The rather sharp transitions in both EGFR^TAIL^ condensation state and Ras activation rate with varying kinase activity level are expected due to the fact that EGFR^TAIL^ condensation occurs via a type of phase transition (2, 4, 31). These results demonstrate that tyrosine kinase inhibition can have the counterintuitive effect of increasing EGFR-mediated Ras activation under certain conditions.

## Discussion and Conclusion

The classical view of EGFR-mediated Ras/MAPK signal activation has been augmented by two recent discoveries: ***i***) EGFR forms a protein condensate involving Grb2 and SOS (2) and ***ii***) SOS activation is strongly dependent on the condensation state of EGFR (2, 5). The further observations that a Grb2:Grb2 interaction through an SH2/SH3 dimer interface contributes to the EGFR condensation process and that Grb2 phosphorylation at Y160 breaks this effect identifies a negative regulatory coupling via Grb2 phosphorylation. Thus, kinase activity that drives EGFR phosphorylation to initiate the signaling process can also inhibit EGFR condensation and Ras activation at higher levels. This dual positive and negative effect of kinase activity is depicted schematically in Figure 7. As experimentally demonstrated here, this can result in counterintuitive effects in which kinase inhibitors effectively promote Ras activation under conditions of high kinase activity.

**Figure 7:**
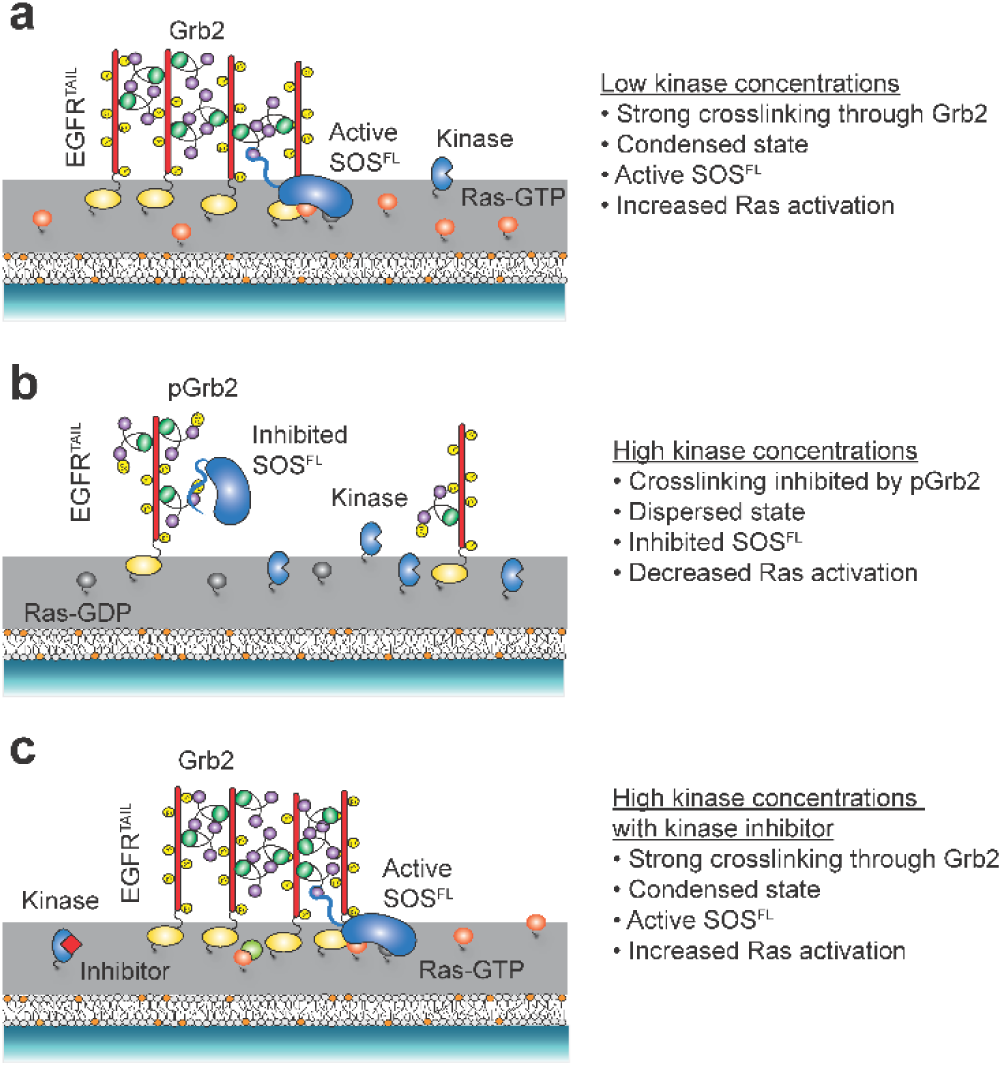
Model of the role of kinases in EGFR condensation. (A) At low kinase concentrations, Grb2 dimerizes and drives EGFR condensation. EGFR condensation promotes SOS^FL^ autoinhibition release and subsequent Ras activation. (B) At high kinase concentrations, Grb2 is phosphorylated and cannot drive EGFR condensation. SOS^FL^ remains autoinhibited, and Ras activation is inhibited. (C) At high kinase concentrations, kinase inhibitors prevent phosphorylation of Grb2. Grb2 can then dimerize to drive EGFR condensation and thereby promote Ras activation.

While the effects of Grb2 phosphorylation on EGFR condensation and Ras activation can be readily reconstituted, it remains unclear the extent to which such effects occur in cells. What is certain is that Grb2 is widely expressed across many cell types and operates under a broad range of healthy physiological conditions (32, 33). Additionally, dysregulation of EGFR signaling through Ras/MAPK is a driver of many human cancers (34, 35). EGFR and Grb2 expression levels vary widely in cancer-derived cell lines (36–39) suggesting that phosphorylation effects on Grb2 and their corresponding effects on Ras activation may occur under certain physiological conditions.

## Author Contributions

H.T.P., C.-W.L., and J.T.G. designed research. H.T.P. and S.J. H. performed research. C.-W.L., J.B.D., and L.J.N.L. contributed new reagents. H.T.P., C.-W.L., and B.L.S. analyzed data. H.T.P., C.-W.L., and J.T.G. wrote the paper.

## Acknowledgements

We thank the members of the Groves Laboratory for feedback on this manuscript. We thank John Kuriyan (University of California, Berkeley) for the helpful discussion and advice. Financial support for this work was provided by National Institutes of Health Grant P01 AI091580 and by the Novo Nordisk Foundation Challenge Programme as part of the Center for Geometrically Engineered Cellular Systems.

## Supporting Citations

References (40–43) appear in the supporting material.

