## Supporting Materials for "Grb2 Phosphorylation Antagonizes EGFR-driven Ras Activation by Interfering with Condensate Assembly"

**Supplemental Information**

**Supplemental Materials and Methods**

**Imaging of SOS Recruitment**

After phosphorylation of EGFR^TAIL^, 750 nM Grb2, 200 pM SOS^FL^-AF55, 0.1 mg/mL BSA, and 1 mM ATP were added into the chamber as a 100 μL injection. After 10 min to establish equilibrium, Images of SOS^FL^-AF555 were acquired with the 561 nm diode laser at 50 mW power and 500 EM gain. Images were collected at a streaming acquisition (shortest possible time between frames) for 1500 frames. Single-molecule images of SOS^FL^ were analyzed using TrackMate (1), an ImageJ plugin, to obtain number of SOS^FL^ recruitment events. SOS^FL^ molecules were localized with Laplacian of Gaussian segmentation, and the initial diameter was set to 3 μm. Tracks were built with a max gap of 2 frames between particle tracks. Quantitative image analysis was done in Python.

**Supplemental Figures and Legends**

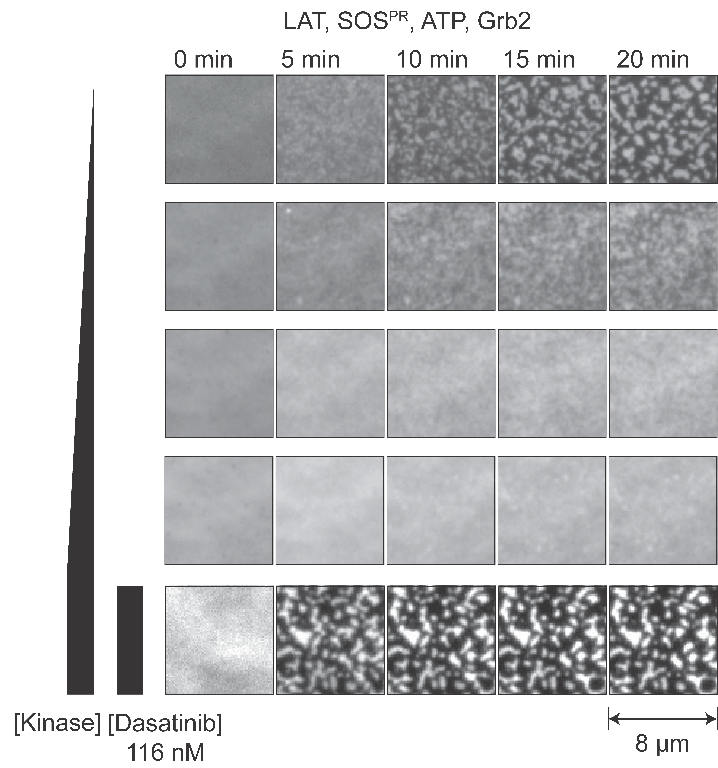

Figure S1: LAT condensation undergoes the same kinase negative feedback regulation as EGFR^TAIL^ condensation. TIRF images of LAT-AF555 reconstituted on SLBs with incubation kinase concentrations of 10 nM, 50 nM, 100 nM, 150 nM, and 150 nM. After addition of Grb2 and SOS^PR^, LAT condenses at low kinase concentrations but remains homogenous at high kinase concentrations. With the addition of kinase inhibitor, LAT forms a condensed phase even at high kinase concentrations (Bottom). Hck was utilized to phosphorylate both LAT and Grb2.

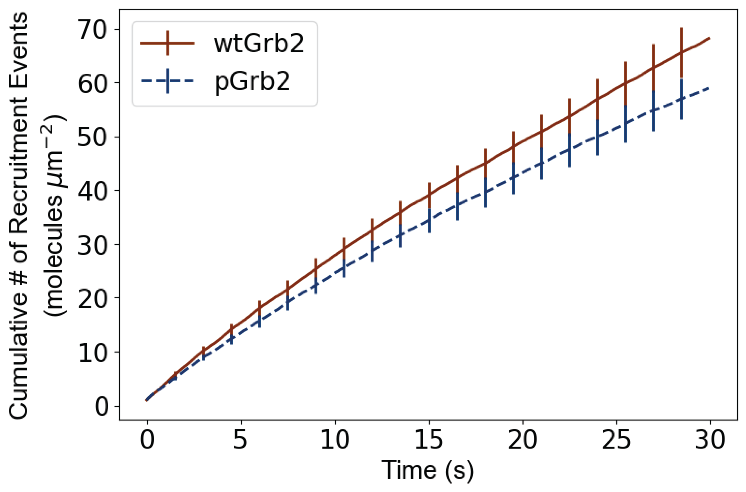

Figure S2: SOS^FL^ recruitment is not dependent on the Grb2 phosphorylation state. Cumulative SOS^FL^-AF555 recruitment events on SLBs functionalized with EGFR^TAIL^-AF488 after the addition of Grb2 (red) or phosphorylated Grb2 (blue). Error bars represent the SEM of 3 replicates.

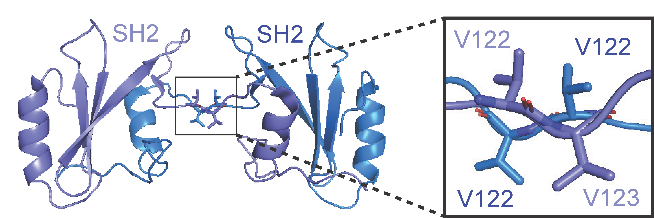

Figure S3: Crystal structure of the Grb2 SH2/SH2 domain-swapped dimer (PDB:6ICH) (2). The position of V122 and V123 is magnified.

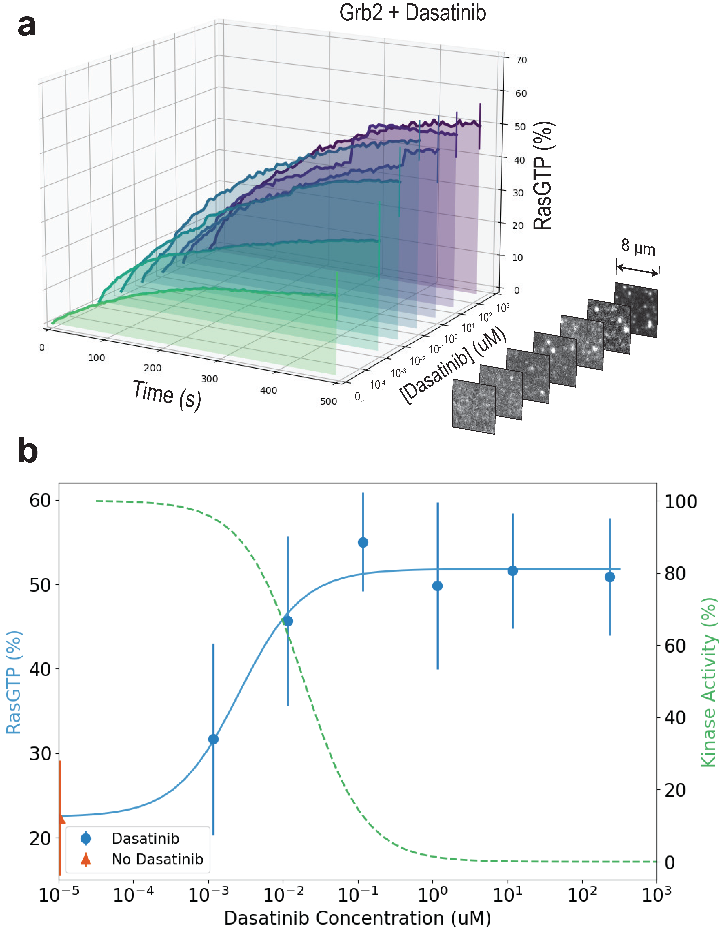

Figure S4: Ras activation is dependent on kinase inhibitor concentration. (A) Mean kinetic traces of ≥ 2 replicates of Ras activation at 150 nM kinase concentration and at different kinase inhibitor concentrations. Error bars represent the SEM of ≥ 2 replicates. Displayed are representative TIRF images of EGFR^TAIL^ on Ras-functionalized bilayers after the addition of Grb2, SOS^PR^, SOS^FL^-AF555, and kinase inhibitor. Increasing kinase inhibitor concentration promotes Ras activation. (B) Dose-response curve of Ras activation as a function of inhibitor concentration. Ras activation increases as kinase inhibitor concentration increases. Error bars represent the SEM of ≥ 2 replicates. A representative Hck activity curve based on the Dasatinib IC_50_ value is overlayed and marked in green (3–5). Ras activation is inversely related to kinase activity.

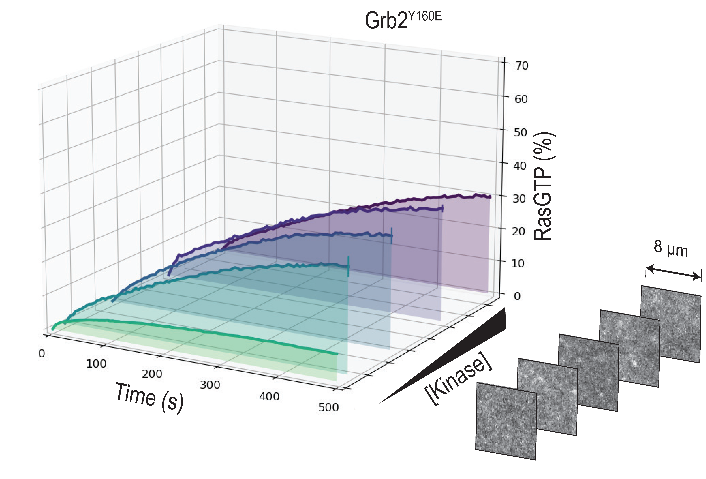

Figure S5: Breaking the Grb2 SH2/SH3 dimer inhibits Ras activation. (A) Mean kinetic traces of ≥ 2 replicates of Ras activation with increasing incubation kinase concentrations of 0 nM, 10 nM, 50 nM, 100 nM, and 150 nM and with mutant Grb2^Y160E^. Error bars represent the SEM of ≥ 2 replicates. Displayed are representative TIRF images of EGFR^TAIL^-AF488 on Ras-functionalized bilayers after the addition of Grb2^Y160E^, SOS^PR^, and SOS^FL^-AF555.

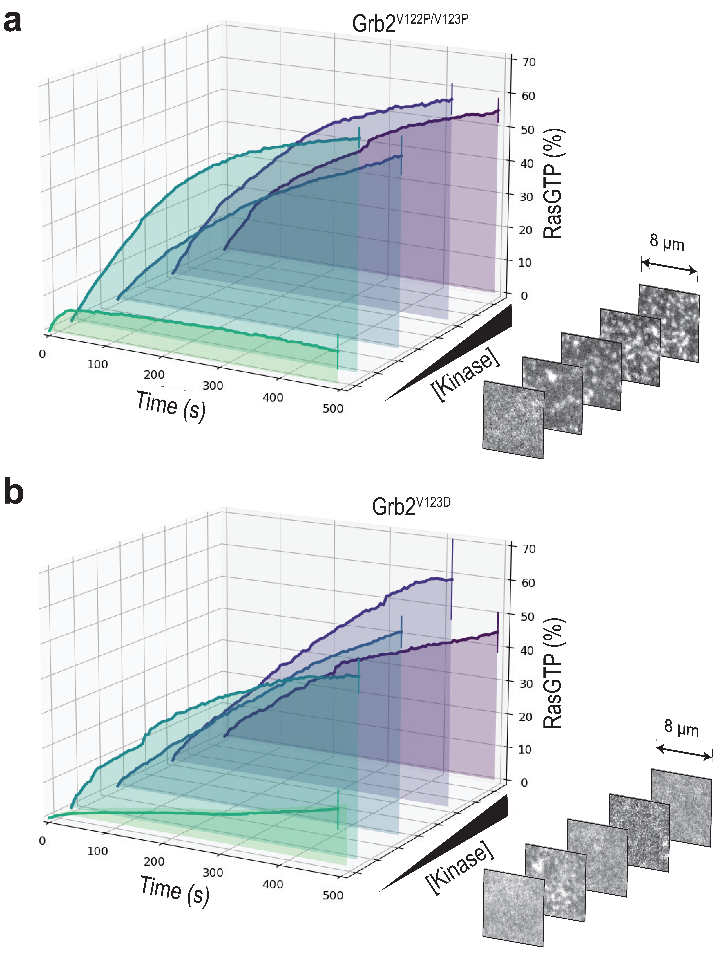

Figure S6: The Grb2 SH2/SH2 domain-swapped dimer can promote Ras activation (A) Mean kinetic traces of 2 replicates of Ras activation at increasing incubation kinase concentrations of 0 nM, 10 nM, 50 nM, 100 nM, and 150 nM and with mutant Grb2^V122P/V123P^. Error bars represent the SEM of 2 replicates. Displayed are representative TIRF images of EGFR^TAIL^-AF488 on Ras-functionalized bilayers after the addition of Grb2^V122P/V123P^, SOS^PR^, and SOS^FL^-AF555. (B) Mean kinetic traces of ≥ 2 replicates of Ras activation at increasing incubation kinase concentrations of 0 nM, 10 nM, 50 nM, 100 nM, and 150 nM and with mutant Grb2^V123D^. Error bars represent the SEM of ≥ 2 replicates. Displayed are representative TIRF images of EGFR^TAIL^-AF488 on Ras-functionalized bilayers after the addition of Grb2, SOS^PR^, and SOS^FL^-AF555. High kinase concentrations inhibit Ras activation and correspond to a dispersed EGFR^TAIL^ state.

**Supplemental Tables**

| Sequence | Modifications | Charge | MH + [Da] | m/z [Da] | Note |
| --- | --- | --- | --- | --- | --- |
| VLNEEcDQNWYK | C6(Carbamidomethyl) | 2 | 1597.69658 | 799.35193 | Tyr 37 |
| VLNEEcDQNWyKAELNGK | C6(Carbamidomethyl);  Y11(Phospho) | 2 | 2289.98516 | 1145.49622 | Tyr 37 |
| VLNEEcDQNWYKAELNGK | C6(Carbamidomethyl) | 3 | 2210.01670 | 737.34375 | Tyr 37 |
| NyIEMKPHPWFFGK | Y2(Phospho) | 3 | 1873.84989 | 625.28815 | Tyr 52 |
| NYIEMKPHPWFFGK |  | 2 | 1793.88445 | 897.44586 | Tyr 52 |
| DIEQVPQQPTyVQALFDFDPQEDGELGFR | Y11(Phospho) | 3 | 3461.55979 | 1154.52478 | Tyr 160 |
| DIEQVPQQPTYVQALFDFDPQEDGELGFRR |  | 3 | 3537.69577 | 1179.90344 | Tyr 160 |
| DIEQVPQQPTYVQALFDFDPQEDGELGFR |  | 3 | 3381.59543 | 1127.87000 | Tyr 160 |

Table S1: Grb2 is phosphorylated by Hck. Peptide analysis of the mass spectrum demonstrates Grb2 is phosphorylated on three different tyrosine residues. Peptide analysis covers 85% of the Grb2 sequence.

5.	Reaction Biology. HCK NanoBRET Target Engagement Intracellular Kinase Assay for screening of compounds binding to the target. *HCK NanoBRET Kinase Assay*.
